# Characterization and identification of a novel daidzein reductase involved in (*S*)-equol biosynthesis in Clostridium C1

**DOI:** 10.1101/2022.01.24.477643

**Authors:** Yun-fei Hu, Chun-Fang Yang, Can Song, Wei-Xuan Zhong, Bai-yuan Li, Lin-yan Cao, Hua-hai Chen, Chang-Hui Zhao, Ye-shi Yin

## Abstract

*(S)*-equol is an isoflavone with high estrogen-like activity and no toxic effects in the human body, and is only produced by some gut bacteria *in vivo*. It plays an important role in maintaining individual health, however, the dearth of resources associated with *(S)*-equol-producing bacteria has seriously restricted the production and application of *(S)*-equol. We report here a novel functional gene C1-07020 that was identified from a chick *(S)*-equol-producing bacterium (Clostridium C1). We found that recombinant protein of C1-07020 possessed similar function to daidzein reductase (DZNR), which can convert daidzein (DZN) into *R/S*-dihydrodaidzein (*R/S*-DHD). Interestingly, C1-07020 can reverse convert *(R/S)*-DHD (DHD oxidases) into DZN even without cofactors or anaerobic conditions. Additionally, high concentrations of *(S)*-equol can directly promote DHD oxidase but inhibit DZNR activity. Molecular docking and site-directed mutagenesis revealed that the amino acid Arg 75 was the active site of DHD oxidases. Subsequently, an engineered *E. coli* strain based on C1-07020 was constructed and showed higher yield of *(S)*-equol than the engineered bacteria from our previous work. Metagenomics analysis and PCR detection surprisingly revealed that C1-07020 and related bacteria may be prevalent in the gut of humans and animals and their *(S)*-equol production state may cause differed between *(S)*-equol producer and non-producer. Overall, a novel DZNR from Clostridium C1 was found and identified in this study, and its bidirectional enzyme activities and wide distribution in the gut of humans and animals provide alternative strategies for revealing the individual regulatory mechanisms of *(S)*-equol-producing bacteria.

**Importance:** *(S)*-equol is a final product of DZN that metabolized by some enteric bacteria. Although *(S)*-equol played very important roles in maintaining human health, larger differences in equol production varied between different populations. Here, a novel DZNR gene C1-07020, which related to *(S)*-equol production, was reported. The bidirectional enzyme functions and wide distribution of C1-07020 in human and animal gut provided additional insights into the metabolic regulation of *(S)*-equol. Additionally, C1-07020 can be used for improving the production of *(S)*-equol in vitro.

## Introduction

In recent years, several long-term follow-up large-scale prospective studies on the intake of soy foods associated with human health, such as those from the USA (1), Western Europe (2), Japan (3, 4) and China (5), have been reported. These have revealed that populations that present eating habits with high isoflavone content from soy products demonstrate an inverse association with cerebrovascular diseases, climacteric period syndrome, and risk of death, in which *(S)*-equol may play an important role. As a soy isoflavone possesses high estrogen-like activity and no toxic side effects, *(S)*-equol, rather than *(R)*-equol, can be an excellent alternative for estrogen in the human body (6). It has become clear that *(S)*-equol is a molecular of interest to researchers and consumers for its various roles in human health. Individuals who are capable of producing *(S)*-equol—as opposed to those who are not—have lower triglyceride and high-density lipoprotein plasma levels and show lower risk rates for cardiovascular diseases (5, 7, 8), For menopausal women, *(S)*-equol producers are at a lower risk of developing osteoporosis (9, 10). Not only that, *(S)*-equol is known to possess several biological activities, such as anti-oxidative stress, anti-inflammatory, and antibacterial activities (10–12), and it can be used for the treatment of atherosclerotic lesions and major symptoms among postmenopausal women (13, 14). Moreover, *(S)*-equol has been used to improve blood glucose homeostasis (15), as well as for the prevention of estrogen-related cancer (16, 17) and osteoporosis (18, 19). Additionally, recent studies have revealed that *(S)*-equol may also have potential utility in patients with Alzheimer disease (20, 21).

*(S)*-equol is not a natural component of soybeans and also cannot be synthesized by the human body. *(S)*-equol is one of the final products of DZN metabolism that occurs in enteric bacteria. Larger differences in equol producers vary between populations, of which there are 20%–60% in Asia, but only about 25% in Western countries (22). Several *(S)*-equol-producing bacteria species have been isolated from human and animal guts. In light of recent findings, the presence of *(S)*-equol-producing bacteria is a *sine qua non* for equol producers (12). Now, *(S)*-equol supplementation has become one of the most effective strategies to improve equol non/low-producer’s health. *(S)*-equol is in great demand for research and biopharmaceuticals, but this demand has to deal with high costs and chiral separation by chemical synthesis, which is currently the main method of isolating *(S)*-equol.

Several studies have focused on *(S)*-equol production in the context of synthetic biology. Depending on study of the five known *(S)*-equol-producing bacteria, *Lactococcus* sp. 20-92 (Lac 20-92) (23), *Slackia isoflavoniconvertens* sp. HE8 (HE8) (24), *Slackia* sp. NATTS (NATTS) (25), *Eggerthella* sp. YY7918 (YY7918) (26) and *Adlercreutzia equolifaciens* sp. DSM19450^T^ (DSM19450^T^) (27), at least four genes have been found to be involved in producing *(S)*-equol, including daidzein reductase (DZNR), dihydrodaidzein racemase (DDRC), dihydrodaidzein reductase (DHDR) and tetrahydrodaidzein racemase (THDR). *(S)*-equol production in these bacteria requires that DZN is first metabolized into DHD by DZNR, and further metabolized into THD and then into *(S)*-equol by DHDR and THDR. Based on the four functional genes of Lac 20–92, we previously constructed a genetically engineered PETDuet-1 DZNR-DDRC PCDFDuet-1 DHDR-THDR BL21 (DE3) (DDDT), which can convert DZN to *(S)*-equol *in vitro* (28). By improving the solubility of DZN, Kim *et al* (29) further improved the production of *(S)*-equol to 1 g/L using a similarly engineered *E. coli* strain. Heterologous synthesis is still the best way for future equol production; however, the improvement of equol production has not yet been explored in the field (30). Continued tapping of new bacteria resources, data mining and furthering a deeper understanding of gene functions, have provided many opportunities for new breakthroughs in *(S)*-equol mass production.

*(S)*-equol-producing bacteria can also be detected in equol non-producers, unfortunately, meaning it is difficult to improve the ability to produce equol, even when isoflavone diet supplements were provided (31–33). A series of studies revealed that the ability of *(S)*-equol production may be associated with equol-related polymicrobial flora and intestinal environment (31, 34). To date, a detailed association between *(S)*-equol-producing bacteria and the ability of individuals to produce *(S)*-equol remain poorly understood. Thus, a key breakthrough for understanding individual differences in *(S)*-equol producer are still focused on the specific bacteria and detailed metabolic mechanics considered. In this study, a chick gut strain (Clostridium C1) was isolated and identified as an *(S)*-equol-producing bacterium, and a novel functional gene involved in DZN metabolism was discussed.

## MATERIALS AND METHODS

### Bacterial culture and *(S)*-equol-producing fermentation

Clostridium C1 was isolated from the caecum of 15-day-old chicks and cultured for 24 h in Reinforced Clostridium Medium (RCM) under anaerobic conditions (10% H_2_, 10% CO_2_, 80% N_2_) at 37°C. Next, these cells were sub-cultured 1:10 in fresh RCM with 40 μM DZN and DHD for 24 h. The fermentation products of C1 were then subjected to analysis by HPLC. First, 1 mL of the fermentation supernatant was mixed with 0.7 mL of ethyl acetate by vigorous shaking and left to sit for 5 min, centrifuged for 5 min at 5 000 g, and the supernatant was transferred to a new centrifuge tube. Each sample was extracted twice following the same procedure. Subsequently, all collected supernatants were dried by vacuum centrifugation and then dissolved in 0.2 ml of methanol. Finally, a Shimadzu HPLC CBM-20A system (Kyoto, Japan) with UV detector SPD-20A was used for HPLC analysis, and separations were performed on a SunFireTM C18 (5 μm, 4.6 × 250 mm) column. Elution was performed using a 0.1% trifluoroacetic acid aqueous solution (A) as well as methanol acetonitrile solution (B, 4:6, V/V), at a flow rate of 0.8 mL/min and a solution B gradient of 50% for 15 min. The detection wavelengths were 205 and 254 nm, the detection temperature was 30°C, and the injection volume was 5 μL.

### Screening for *(s)*-equol-producing functional genes

Clostridium C1 was grown in 100 ml of Gifu anaerobic medium (GAM) for 24 h, and after centrifugation at 5 000 g for 5 min, bacteria were collected and genomic DNA was extracted using a Gram-positive bacterial DNA extraction kit (Qiagen, Germany). This genomic DNA was subjected to whole-genome sequencing by next-generation sequencing (Nextomics, China). A whole-genome reference for C1 was compared with reported DZNRs (Lac 20–92, Accession number AB558141.1, DSM19450, Accession number AP013105.1, HE8, Accession number JQ358709.1, NATTS, Accession number AB646272.1, YY7918, Accession number AP012211.1) using Local BLAST (https://blast.ncbi.nlm.nih.gov/Blast.cgi) analysis. Potential functional genes were searched against the NCBI conserved domain database (CDD, https://www.ncbi.nlm.nih.gov/Structure/cdd/wrpsb.cgi), then the obtained *(S)*-equol-producing functional gene clusters were analyzed using Motif Scan (https://myhits.sib.swiss/cgi-bin/motif_scan). Likewise, other novel functional genes were screened using the same method for reported DDRCs, DHDRs, and THDRs.

### Reductase activity detection of C1-07020 in vitro

The C1-07020 gene was cloned into the plasmid pETDuet-1 (Novagen) and then transformed into *E. coli* BL21 (DE3) (forward primer: 5’-ccc<u>gcggccgc</u>atgaaaaacaaatattaccctca-3’, reverse primer: 5’-ccg<u>cctagg</u>ttagatttgtctggctgctat-3’, restriction enzyme sites are underlined). After induction and expression in LB liquid medium, C1-07020 protein was purified by Ni^2+^ affinity chromatography, and protein purity and concentration were determined by SDS-PAGE and BCA protein assay (Solebro), respectively. The DZNR activity of purified C1-07020 was detected in 1 ml of potassium phosphate buffer (100 mM, pH 7.0) under anaerobic conditions. 3 μM C1-07020 recombinant protease, 80 μM DZN, 500 μg/ml of NADPH/NADH, 0.5 mg/ml of sodium bisulfite, 0.3 mg/ml of dithiothreitol, and 0.2 mg/ml phenylmethylsulfonyl fluoride were used. Following incubation at 37°C for 4 h, samples were extracted by ethyl acetate, and then used for HPLC detection. Elution was performed under 30% acetonitrile with a 0.1% acetic acid solution and a flow rate of 1 mL/min for 15 min. The detection wavelengths were 205 and 275 nm, the detection temperature was 30°C, and the injection volume was 10 μL. Moreover, the S-enantiomer and R-enantiomer of DHD were detected using to a previously published procedure (23). To further characterize the role of equol-production, C1-07020 was constructed in an *(S)*-equol producing engineered *E. coli* DDDT established in our preliminary work (28), but instead with DZNR replaced by C1-07020 (C1-07020DDT). Fermented productions were compared between DDDT and C1-07020DDT using our previously reported method (28).

### Oxidases activity assay of C1-07020

To identify the novel oxidase function of C1-07020, the DZNRs from Lac 20-92 (GenBank: BAJ22678) and NATTS (GenBank: BAL46930) were selected as controls. Both the gene sequences of DZNRs were synthesized (Genscript, China) and cloned in pETDuet-1 and then transformed into *E. coli* using a similar method as with C1-07020. pETDuet-1 Lac 20-92 DZNR BL21 (DE3) and pETDuet-1 NATTS-DZNR BL21 (DE3) were cultured in LB medium; then each of the recombinant proteins was induced and purified. Subsequently, recombinant proteins for C1-07020, DZNRs from Lac 20–92 and NATTS were used to identify DHD oxidase function in 1 ml of 0.1 M PBS (pH7.0) with 80 μM of DZN or DHD (the same method of reductase activity detection as mentioned previously).

DHD oxidase activity using C1-07020 recombinant protein was next detected *in vitro*. First, the optimal pH was determined in 1 ml 0.1 M citrate buffer (pH 5.0, 5.5, 6.0, or 6.5) or 0.1 M PBS (pH 7.0, 7.5, or 8.0) with 80 μM *(R)*-DHD or *(S)*-DHD and 3 μM recombinant protein from C1-07020. Samples were incubated at 37°C for 2 h and then DZN was extracted for detection by HPLC. The temperatures for detection were 30°C, 34°C, 37°C, 41°C, and 45°C. After that, k enzyme reactions were carried out for one hour. To determine the oxygen sensitivity, 0.5 mg/ml cysteine hydrochloride was added according to optimal conditions above and used for enzyme activity detection under both aerobic and anaerobic conditions. In order to evaluate the influence of reductase and oxidases activity, *(S)*-equol (Daicel, China) and estradiol (Solebro) at concentrations of 0.08, 0.16, 0.32, 0.64, 1.28, 2.56, and 4 mM were added to DZN or DHD enzyme reaction mixes using C1-07020 recombinant protein, and then evaluated by HPLC.

### Molecular docking study

Molecular docking was used to investigate the key binding mode between the compound and C1-07020 using Autodock vina 1.1.2 (35). The three-dimensional (3D) structure of C1-07020 was built using SWISS-MODEL, a fully automated protein structure homology-modeling server. The 2D structure of the compound was drawn using ChemBioDraw Ultra 14.0 and converted to a 3D structure using ChemBio3D Ultra 14.0 software. The AutoDock Tools 1.5.6 package (36) was employed to generate docking input files. The ligand was prepared for docking by merging non-polar hydrogen atoms and defining rotatable bonds. The search grid for the C1-07020 site for DHD was identified as center_x: −18.109, center_y: −4.869, and center_z: 6.716 with dimensions size_x: 21.75, size_y: 15, and size_z: 15. The search grid of the C1-07020 site for DZN was identified as center_x: −18.109, center_y: −4.869, and center_z: 6.716 with dimensions size_x: 21.75, size_y: 15, and size_z: 15. The search grid of the C1-07020 site for NADH was identified as center_x: −27.531, center_y: −8.022, and center_z: −1.784 with dimensions size_x: 15, size_y: 15, and size_z: 15.75. In order to increase the docking accuracy, the value of exhaustiveness was set to 20. For Vina docking, the default parameters were used if not otherwise mentioned. The best-scoring pose as judged by the Vina docking score was chosen and visually analyzed using PyMoL 1.7.6 software (www.pymol.org). Meanwhile, SMART tool was used for function prediction (http://smart.embl-heidelberg.de/).

### Site-directed mutagenesis

Based on the molecular docking prediction, eight potential enzyme functional regions in C1-07020 were selected for multiple amino acid mutation (Table S1). Briefly, two overlapping fragments for each mutation were amplified using fp1-rp1 and fp2-rp2, and overlap extension PCR with fp1-rp2 was performed for fusing full-length gene of mutants. The obtained mutants were cloned into pETDuet-1 using a one-step PCR kit (Vazyme, China). Subsequently, recombinant plasmids were transformed into *E. coli* BL21 (DE3). Positive transformers were identified by PCR and then confirmed by Sanger sequencing (Sangon, China). Recombinant proteins from mutants were induced as small scale clones of C1-07020, and enzyme function was determined from 1-ml whole-cell biocatalyst with 0.2 mM of DZN or DHD added, followed by incubation at 37°C for 4 h. Samples were then extracted and assessed by HPLC. Target mutants were selected for purification and used for quantitative detection with the same method as our activity assays for C1-07020.

Following the results of multiple amino acid mutation, 16-point mutations were selected and used for identifying the key amino acid active site (Table S2), and each point mutant was analyzed by using the above method. For further evaluating the key functional region of DHD oxidases activity, another site-directed mutagenesis screen was performed on the DZNR from Lac 20-92. The DZNR gene was used as a target, and amino acids at positions 72-78 were replaced by C1-07020 with the key amino acid site of DHD oxidases activity (Table S3).

### Metagenomics analysis of C1-07020 in the human gut

To assess the distribution of these DZNR genes in the human intestine, macrogenomics analysis was applied. Briefly, 4644 reference genomes were downloaded from the NCBI database of the human gut microbiome (37) to test for similarity with C1-07020 protein sequence using the tblastn tool, and > 40% matching similarity and > 90% coverage were used as the selective parameters. Second, based on the obtained reference genomes with high similarity and coverage greater than 80%, genes with the same types as Clostridium C1 were screened and downloaded from NCBI and tested against the C1-07020 protein sequence using the above method. Afterward, all genes obtained from our two-step macrogenomic analysis, as well as the seven reported DZNR genes (Lac 20-92: BAJ22678, YY7918: WP_013979957, HE8: WP_123220034, NATTS: BAL46930, DSM19450: WP_022741749, AUH-JLC159: AIC80887, Slackia: WP_123208882), were aligned using Evolutionary Genetic Analysis (MEGA) (38).

One gene (00242_GENOMEO01471_13) with about 80% similarity and two genes (GUT_GENOME103990_60, GUT_GENOME127776_11) (37) with about 40% similarity for C1-07020 were sequence synthesized and functional testing of these genes was performed as described for C1-07020.

### Analysis of sequence diversity

Based on the results of macrogenomics analysis, primers for cluster 1 and cluster 2 of these DZNRs were designed for PCR detection from the feces of humans and mice (Table 1). Briefly, 30 fecal samples were collected from 6–8-week-old female mice from Hunan SJA laboratory animal CO., LTD (China), and another 20 fecal samples were collected from 20–30-year-old healthy female volunteers. Samples ranged from 0.2 to 0.3 g per sample, which were then resuspended in water and the supernatants were obtained by centrifugation at 5,000 g for 10 min to detect *(S)*-equol as described above. The remaining samples were then used for total DNA extraction using a fecal DNA Kit (QIAGEN, Germany). PCR was performed to investigate the distribution of DZNRs, *E. coli* BL21 selected as a negative control.

**TABLE 1.**
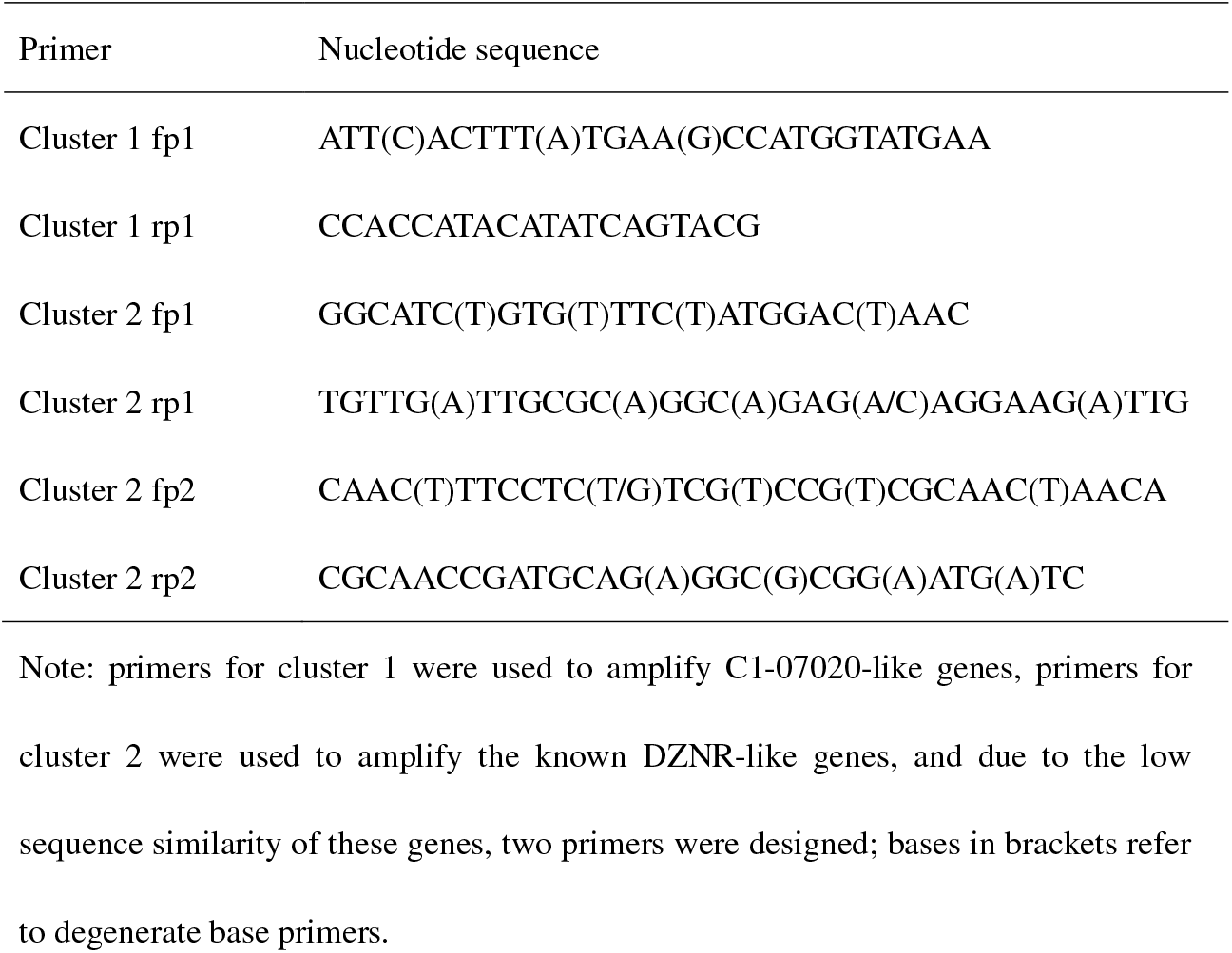
Primers used for detection of C1-07020 and DZNRs

### Statistical analysis

Each treatment or group was assessed at least three times in parallel, and all data was analyzed using GraphPad Prism8 (San Diego, CA). The results for all data are displayed as the mean and standard deviation (SD). Statistical significance for DZN, DHD and *(S)*-equol were determined using Two-way ANOVA followed by a Tukey–Kramer comparisons *post hoc* test. Oxidases activity assay of C1-07020 followed Michaelis–Menten Kinetics. In addition, statistical differences in the enzyme reaction products were analyzed by Student’s *t*-test and the significance level was set at P < 0.05.

## RESULTS

### Clostridium C1 *(s)*-equol-production and functional genetic screen

Clostridium C1, which is a Gram-positive, filamentous, anaerobe bacterium isolated from chick cecum, can transform DZN into DHD and *(S)*-equol under strictly anaerobic conditions (Fig S1). However, when DHD instead of DZN was used as substrate, DZN was also detected in the fermentation broth of Clostridium C1. The complete genomic sequence of Clostridium C1 was obtained (GenBank CP073631) and used for an *(S)*-equol-producing functional genetic screen. After comparing sequence similarity and functional domains with the five reported DZNRs, a potential target gene in C1-07020 was found and selected for enzyme functional assays *in vitro* (Fig S2).

### Enzyme activity analysis of C1-07020

C1-07020 was cloned into pETDuet-1 and expressed in *E. coli* BL21 (DE3), and soluble recombinant proteins were purified and used for DZN transformation assays. The results showed that recombinant protein from C1-07020 possessed DZNR activity that could transform DZN to DHD (Fig S3). Interestingly, both *(S)*-DHD and *(R)*-DHD were detected by chiral HPLC analysis, and the 68.9% of the total product was *(S)*-DHD (Fig 1-A). Furthermore, as with the reported DZNRs, C1-07020 was an NADH/NADPH-dependent dehydrogenase that required anaerobic conditions. To further evaluate the impact of C1-07020 in *(S)*-equol-producing, Lac 20–92 DZNR from the strain DDDT was replaced with C1-07020 and used to construct C1-2079DDT. The fermentation results are shown in Fig. 1-B, and C1-07020DDT could produce *(S)*-equol at about 10% higher levels than DDDT at a substrate concentration of 0.16 mM (p < 0.05), although it requires a longer fermentation time (8 h) than DDDT (2 h).

**FIG 1.**
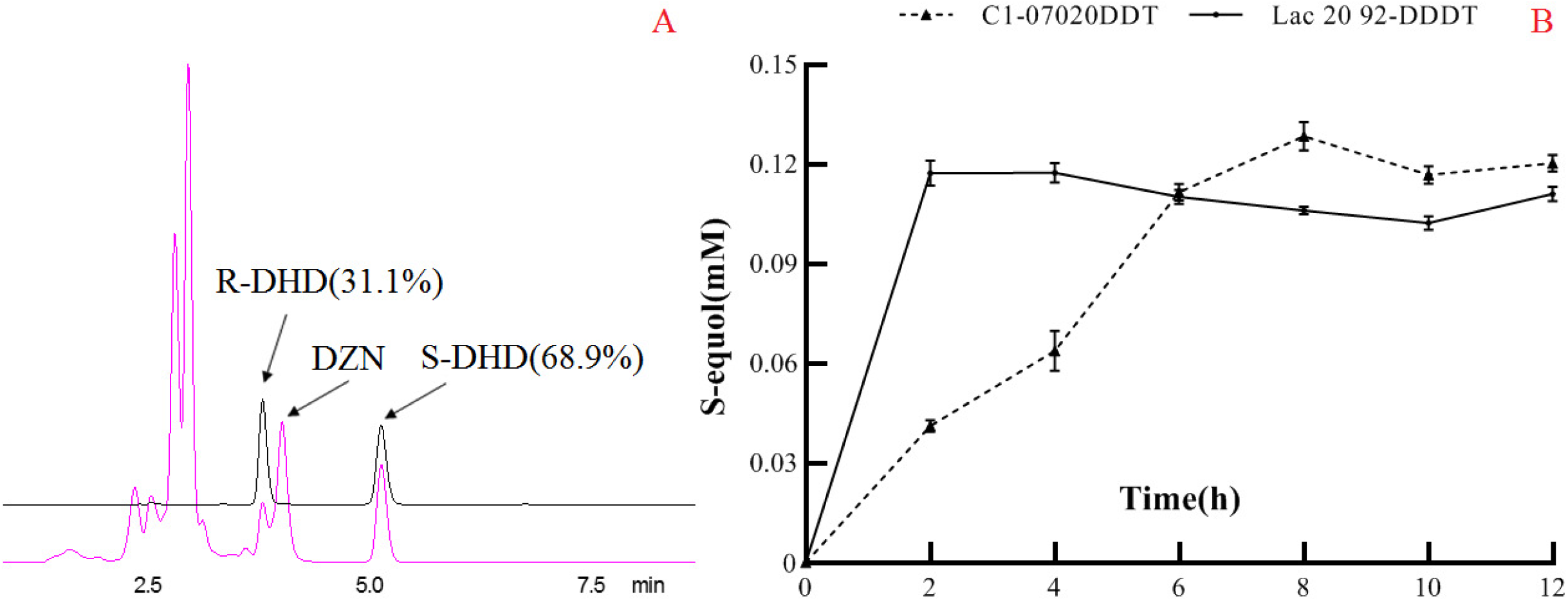
Reductase enzyme activity assays using C1-07020. A: chiral detection of DHD by HPLC; B: comparative analysis of *(S)*-equol production between C1-07020DDT and Lac 20-92 DDDT.

In order to probe the DHD reverse transformation function of C1-07020, DZNRs from Lac 20–92 and NATTS were selected as controls. The results of enzymatic assays indicated that all recombinant proteins showed DZNR activity, which is an ability to convert DZN to DHD, but only C1-07020 demonstrated an oxidase activity, which is converting DHD to DZN (Fig. 2A). Further oxidases activity analysis showed that the optimal conditions for C1-07020 were a pH of 6.5 and a temperature of 41°C, and kinetic parameters for this reaction are *V_max_* of 52.2 and a *K_m_* of 4.237. It was also observed that the optimal time for the enzyme–substrate reaction was 6 h at a DZN conversion rate of 57.3% when 80 μM DHD was added (Fig. 2 B-D). Both *(R)*-DHD and *(S)*-DHD could be converted into DZN by C1-07020, and did not require additional coenzymes or anaerobic conditions.

**FIG 2.**
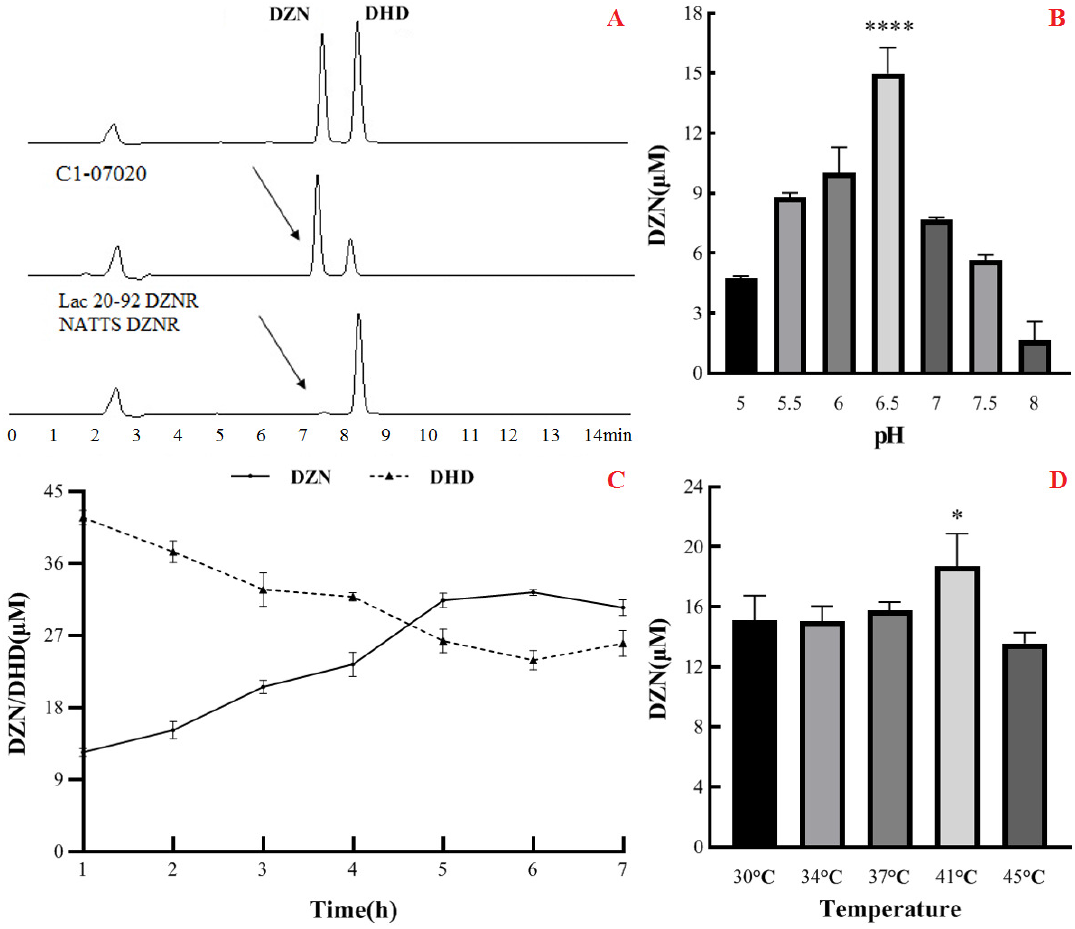
DHD oxidase activity with C1-07020. A: The DHD reverse transformation function was detected by HPLC; B: the optimal pH for oxidase activity; C: kinetic analysis of oxidase activity; D: the optimal temperature for oxidase activity. The results of enzymatic assays indicated that all recombinant proteins showed DZNR activity, which is an ability to convert DZN to DHD, but only C1-07020 demonstrated an oxidase activity, which is converting DHD to DZN. Further oxidases activity analysis showed that the optimal conditions for C1-07020 were a pH of 6.5 and a temperature of 41 °C, and kinetic parameters for this reaction are *V_max_* of 52.2 and a *K_m_* of 4.237. It was also observed that the optimal time for the enzyme-substrate reaction was 6 h at a DZN conversion rate of 57.3% when 80 μM DHD was added (Figure 2 B-D). Both *(R)*-DHD and *(S)*-DHD could be converted into DZN by C1-07020, and did not require additional coenzymes or anaerobic conditions.

### Sequence analysis and molecular docking

The gene cluster for *(S)*-equol-producing enzymes was next considered, as it is very similar for all reported *(S)*-equol-producing bacteria, but not for genes similar to DDRCs, DHDRs and THDRs, as these have not been found in Clostridium C1 (Fig. S2). Moreover, 12-kb upstream and downstream of C1-07020 was identified and no functional genes were identified (Fig. S4). Furthermore, analysis showed that C1-07020 showed conserved domain similarity to DZNRs (Fig. S5) that also have an OYE-like FMN binding domain (potential substrate binding or enzyme activity site), a 4Fe-4S cluster motif or coenzyme binding motifs (potential enzyme activity site).

To further explore the novel oxidases function of C1-07020, a molecular docking study was used to screen this enzymatic activity site. DHD was docked into the binding site of C1-07020 and the results are shown in Fig. 3A. This study revealed that DHD was located to the hydrophobic site, surrounded by the residues Pro-106, Ala-173, Ile-349 and Met-353, forming a stable hydrophobic structure. Detailed analysis showed that the chromone scaffold of the DHD formed CH-π interactions with the residue Tyr-32 and Cys-72. Importantly, two key hydrogen bond interactions were observed between the DHD and the residues His-261 and Ile-349, which was the main interaction between the DHD and C1-07020. For the reductase activity of C1-07020, DZN and NADH were docked into the binding site of C1-07020 (Fig. 3B). The results showed that the residues Pro-106, Arg-108, Phe-134, Gly-174, His-261, Ile-349, and Met-353 may be the potential function cites of C1-07020.

**FIG 3.**
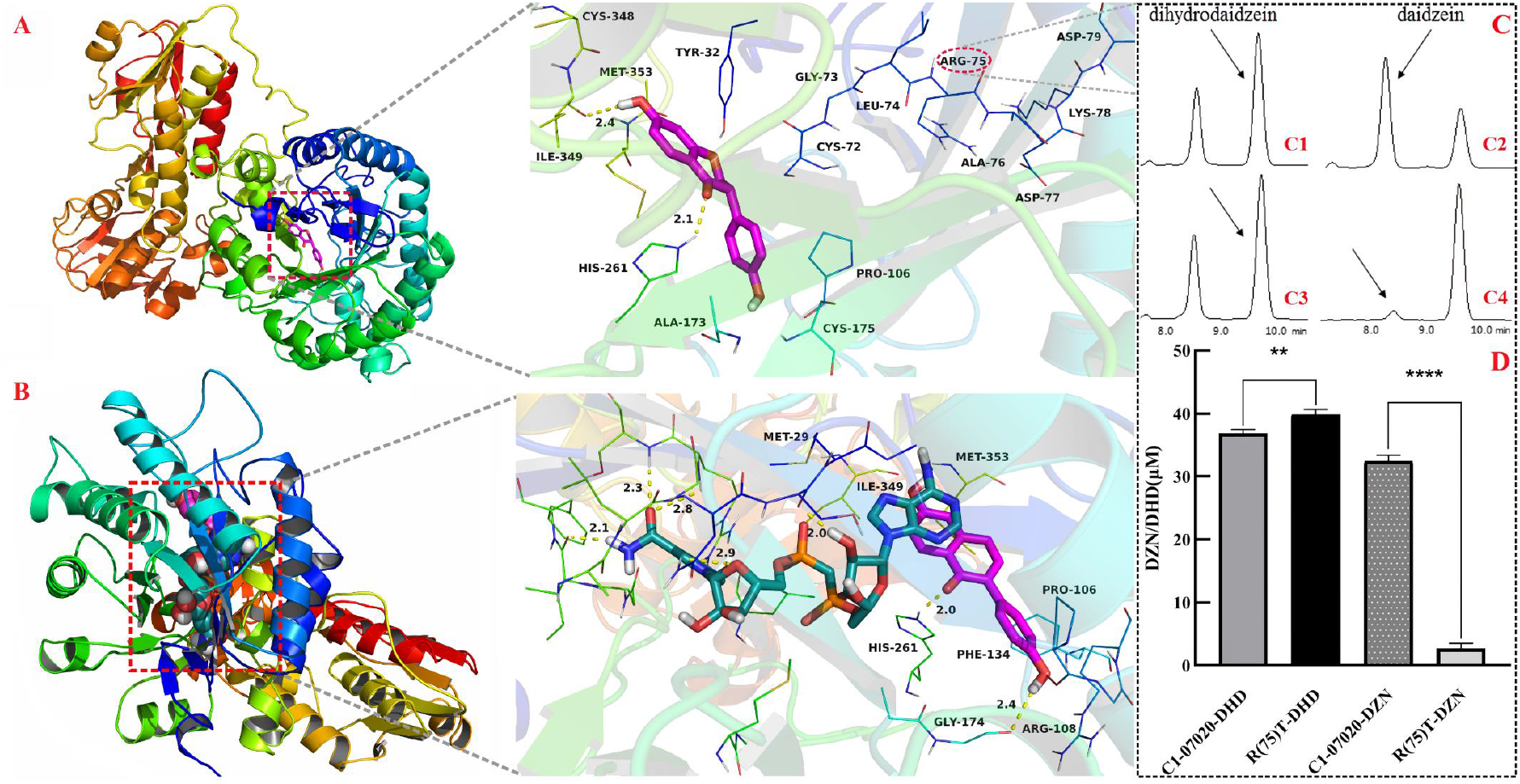
Analysis of oxidase key functional loci in C1-07020. A: DHD was docked into the binding site of C1-07020; B: DZN and NADH were docked into the binding site of C1-07020; C: function detection of R(75)T C1-07020 mutation by HPLC. C1 and C2 respectively denote reductase and oxidase functions of C1-07020; C3 and C4 respectively denote reductase and oxidase functions of R(75)T C1-07020 mutation; D: quantitative detection of C1-07020 and R(75)T C1-07020 mutation, DHD and DZN denote reductase and oxidase function detection, respectively. Molecular docking revealed that the residues Pro-106, Arg-108, Phe-134, Ala-173, Gly-174, Ile-349, Met-353, Tyr-32, Cys-72, His-261 and Ile-349 may be the potential function cites of C1-07020, these residue sites were selected to site-directed mutagenesis and the results showed that amino acid at position 75 Arg was identified as a key amino acid with DHD oxidases function of C1-07020.

### DHD oxidases function site assays

Following the results of molecular docking predication, eight potential functional regions in C1-07020 were examined by site-directed mutagenesis with mutations consisting of several either consecutive or inconsecutive amino acid substitutions (Table S1). We found that DHD oxidase activity was significantly weakened with mutagenesis of the amino acid sequence at positions 72–74 and 79 (72<u>CGL</u>RADK<u>D</u>79/AKHRADKA), but the DZN reductase activity was also weakened to some extent. A further mutagenesis based on amino acid sequence at positions 75–78 (75<u>RADK</u>78/TTFI) revealed that the DHD oxidase activity of mutant 75–78 was almost lost; however, there was no significant change in DZN reductase activity. Other mutants either showed no change or were lost, or decreased significantly all enzyme activity (Table S4).

Point mutations at positions 75–78 were then further analyzed. The results from quantitative examination showed that the R(75)T mutant lost essentially all DHD oxidases function, but the corresponding DZN reductase activity was increased (p < 0.01). Finally, the amino acid at position 75 (Arg) was identified as the key amino acid for DHD oxidase function in C1-07020 (Figure 3C-D). Moreover, site-directed mutagenesis of A(76)T, D(77)F, K(78)I and D(79)A resulted in reduced DHD oxidase activity to varying degrees and no significant change in DZN reductase activity. Other point mutations at positions 72–74 were also assessed, and the enzyme activities of these mutants were either not significantly different or decreased to some extent for both DZN and DHD (Table S4).

After identifying the key active site of DHD oxidase function in C1-07020, directed evolution of Lac 20–92 DZNR was performed by swapping in the active sites of C1-07020. We found that a DZNR mutant with S(75)R resulted in DHD oxidase activity (Fig. S6).

### The effect of *(s)*-equol and estradiol on enzyme activity

The efficiency of Clostridium C1 for producing *(S)*-equol is affected by the product concentration, due to the reversible enzyme activity of C1-07020, the product of *(S)*-equol. The structurally similar compound estradiol was applied to these experiments, and the regulatory effects on enzyme functions were subsequently evaluated. When the concentration of the fermented DZN was 80 μM, the addition of *(S)*-equol at concentrations ranging from 20 to 320 μM had no effect on enzyme activity (p > 0.05).

However, with a concentration of *(S)*-equol exceeding 640 μM, the reaction proceeded toward DZN formation and showed a concentration dependent increase in oxidase activity with increasing *(S)*-equol concentrations (p < 0.001). In contrast, even when estradiol was added to a saturating concentration (4 mM), there were no significant effects (Fig. 4). Moreover, fermentation of DHD was also evaluated with the addition of *(S)*-equol or estradiol using the same method, and similar results were also observed.

**FIG 4.**
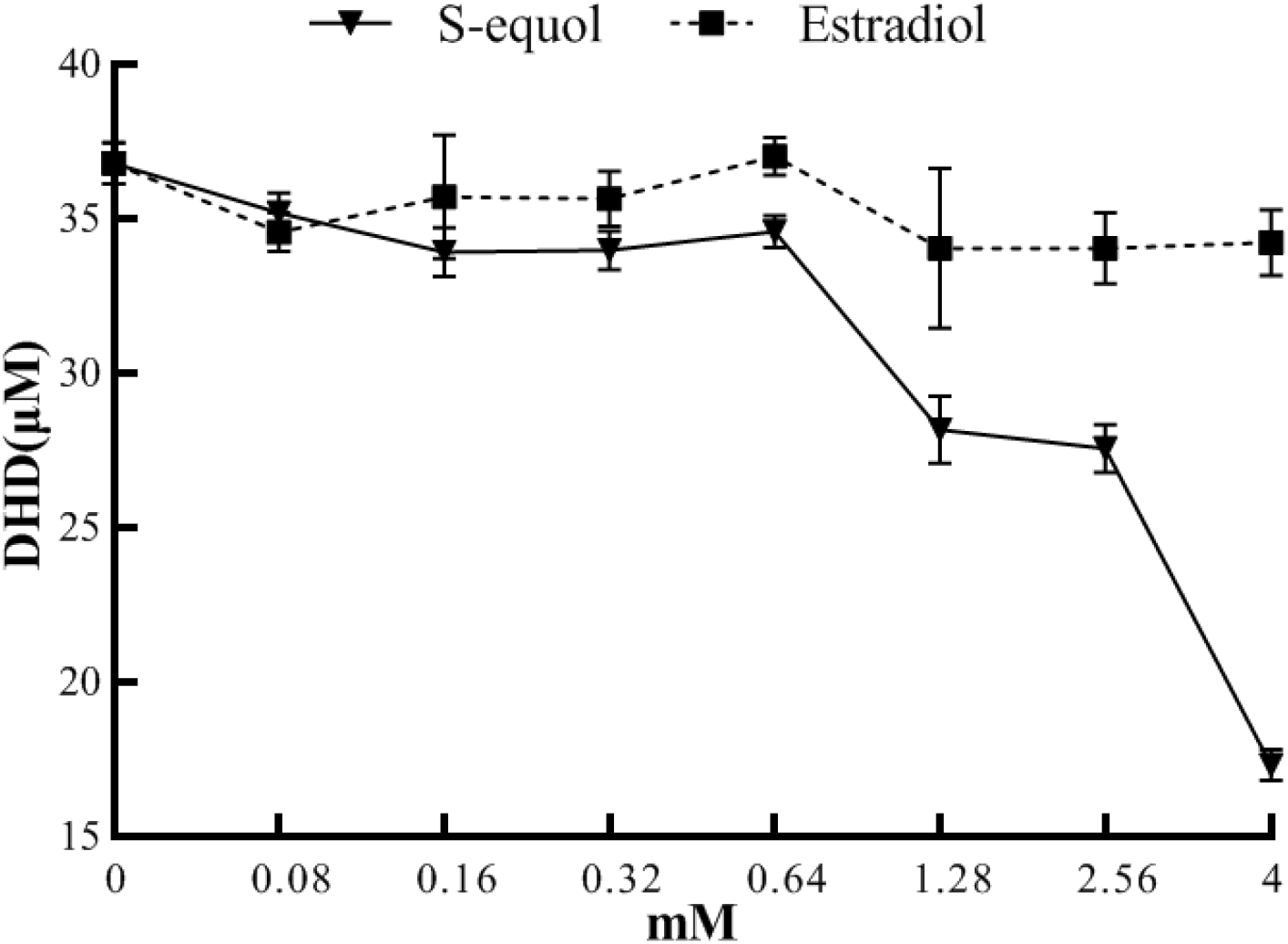
Effect of *(S)*-equol or estradiol concentration on reductase and oxidase activity with C1-07020. With a concentration of *(S)*-equol exceeding 640 μM, the reaction proceeded toward DZN formation and showed a concentration dependent increase in oxidase activity with increasing *(S)*-equol concentrations (p < 0.001). In contrast, even when estradiol was added to a saturating concentration (4 mM), there were no significant effects.

### Metagenomics analysis of C1-07020

The amino acid sequence of C1-07020 showed 100% identity with a protein (GenBank: OUQ08445) that had been metagenomic sequenced from chicken caecum. In this study, C1-07020 was identified as a novel DZNR with both DZN reductase and DHD oxidases activities. To further explore the biological significance of this gene, C1-07020 was used for amino acid sequence alignment analysis against the NCBI human gut metagenome database. Twenty-nine genes showed high similarity at the amino acid sequence level with C1-07020, of which, the three reference strains MGYG-HGUT-00242 (containing 880 genomes), MGYG-HGUT-00244 (containing 692 genomes), and MGYG-HGUT-02689 (containing 33 genomes), had coverage greater than 80% for C1-07020. Genes in the three reference strains were then used to blast C1-07020 again and another twenty sequences were obtained. Following evolutionary genetic analysis, three clusters with C1-07020 could be distributed in an evolutionary tree (Fig. 5). Twenty-three genes showed a high similarity with C1-07020 in the range from 79–100% and are listed in cluster 1, and almost half of these genes shared more than 90% similarity, particularly, 00242_GENOME214702_93, which was completely consistent with C1-07020 and this sequence had only one amino acid difference. In addition, cluster 2 was shown to have similarity with C1-07020 between 44–46%, and most of these genes have been identified and reported as DZNRs.

**FIG 5.**
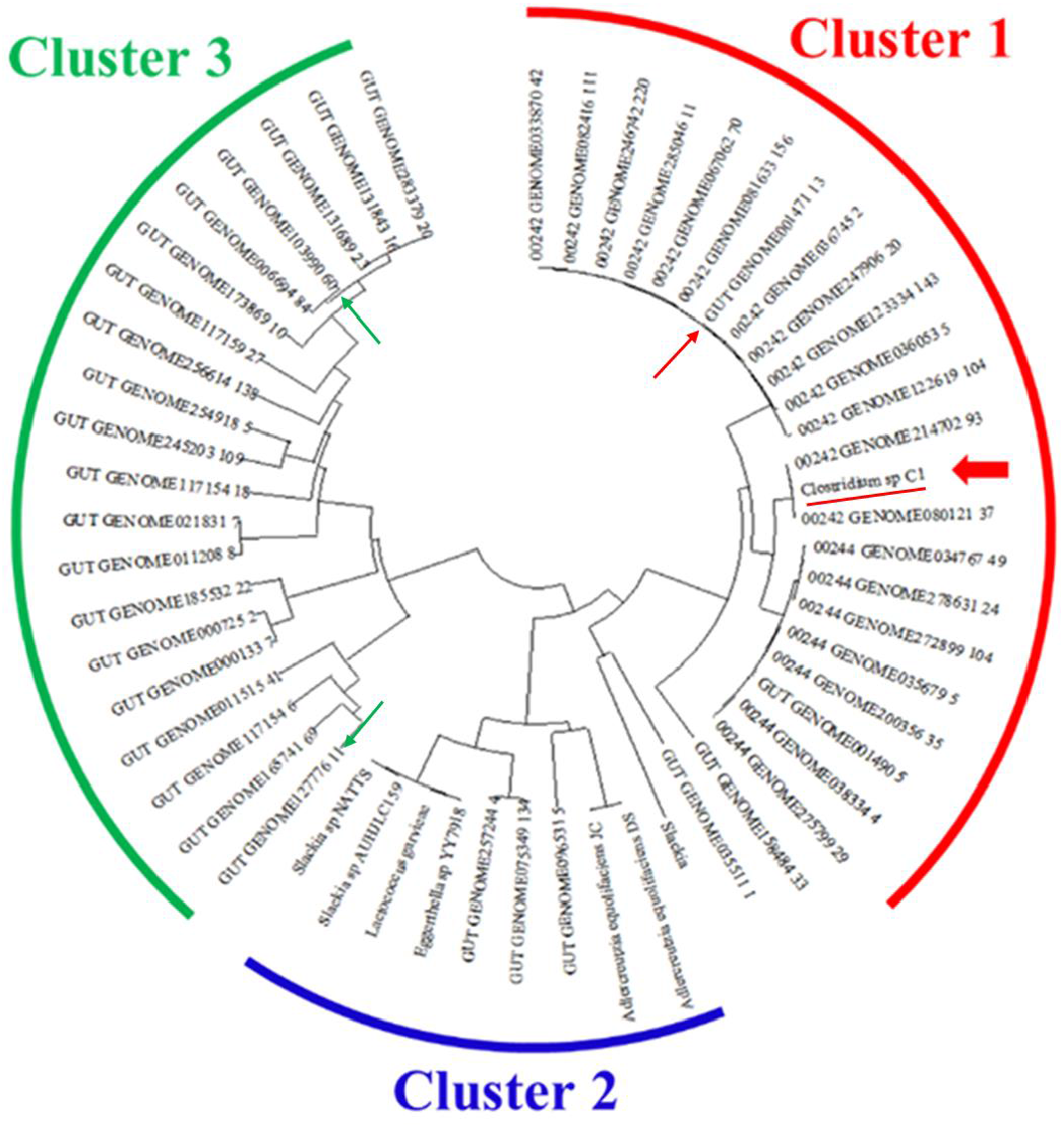
Bioinformatics analysis of C1-07020 across the human gut metagenome. Cluster 1: the similarity of amino acid sequences with C1-07020 were between 79-100%; Cluster 2: the similarity of amino acid sequences with C1-07020 were between about 44-46%; Cluster 3: the similarity of amino acid sequences with C1-07020 were between about 40%. C1-07020 is marked by underline, two genes showed low sequence similarity (marked by green arrows) and one gene showed high sequence similarity (marked by red arrow) with C1-07020 were selected to functional identify *in vitro*.

In order to validate these predictions, two genes that showed about 40% sequence similarity in cluster 3 and one gene that showed 80% sequence similarity from cluster 1 with C1-07020 were selected for functional identification *in vitro*. As expected from bioinformatics analysis, the genes in cluster 1 had DZN reductase and DHD oxidase activity (Fig. 6A), However, no DZNRs were found in cluster 3 (Fig. 6B). To further investigate the distribution of C1-07020-like genes, *(S)*-equol and PCR were used to detect these in the feces of humans and mice. The detection rates of both *(S)*-equol and C1-07020-like genes were almost 100% in mice, and about 40% of the genes in cluster 2 were also found. However, about 75% of fecal samples from humans contained C1-07020-like genes, but only a small percentage of these were positive for cluster 2 genes and *(S)*-equol (Fig. 6C).

**FIG 6.**
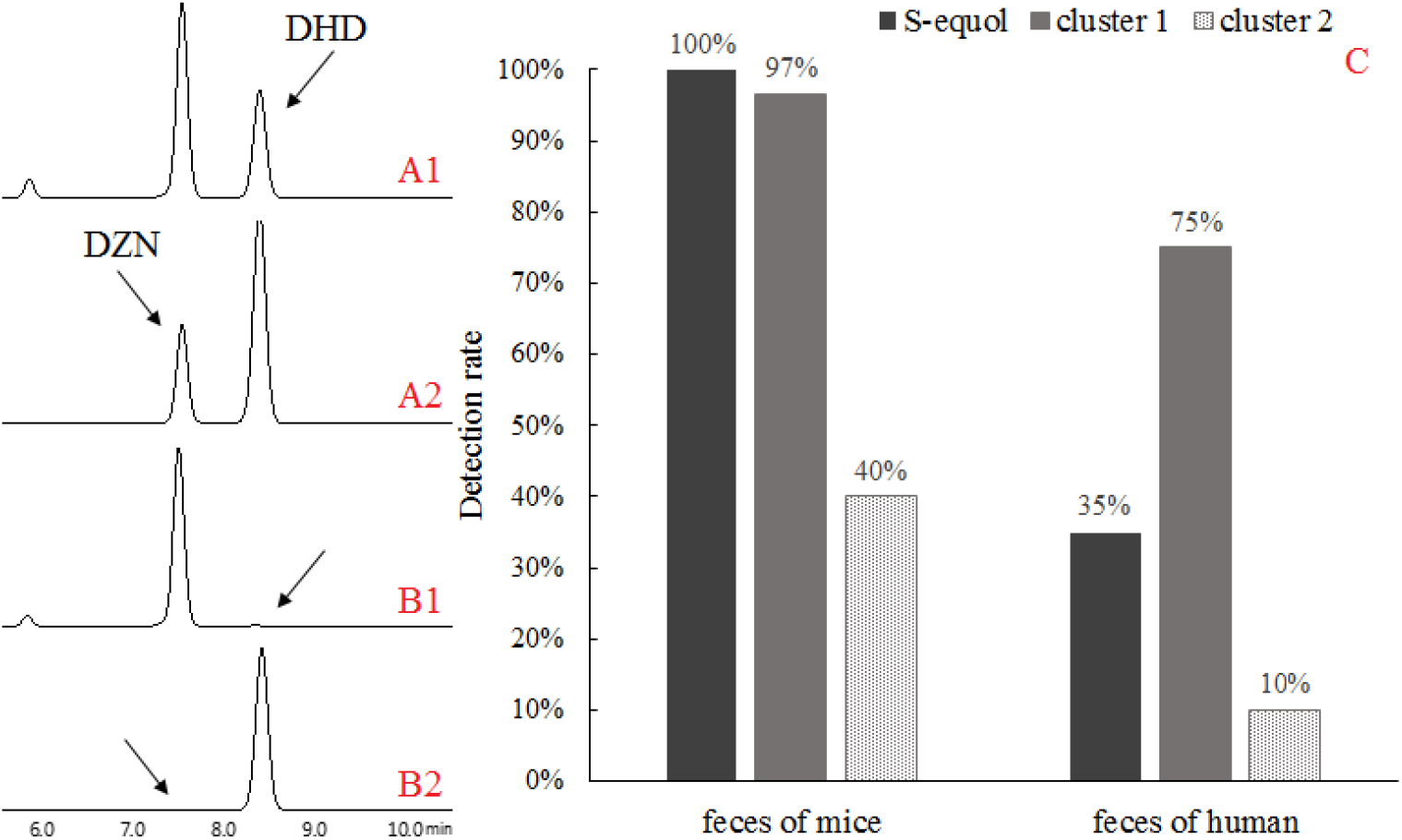
Detection of *(S)*-equol and C1-07020-like genes in human and mice feces. A1 and A2 denote DZN reductase and DHD oxidase function from 00242_GENOMEO01471_13, it showed that the genes in cluster 1 had DZN reductase and DHD oxidase activity; B1 and B2 denote DZN reductase and DHD oxidase function from GUT_GENOME103990_60 and GUT_GENOME127776_11, however, no DZNRs were found in cluster 3; C: detection rate of *(s)*-equol (HPLC) and C1-07020-like genes (PCR) in human and mice feces. The detection rates of both *(s)*-equol and C1-07020-like genes were almost 100% in mice, and about 40% of the genes in cluster 2 were also found. However, about 75% of fecal samples from humans contained C1-07020-like genes, but only a small percentage of these were positive for cluster 2 genes and *(s)*-equol.

## DISCUSSION

The strain Clostridium C1 was isolated from chick gut. In anaerobic fermentation, DZN was metabolized to DHD and *(S)*-equol by this strain, to our surprise, when DHD was used as the substrate, and DZN was also detected in the fermentation products. It is well known that DHDR and THDR can catalyze a reversible reaction, but not DZNR (23-27). Therefore, this is the first discovery of the ability of *(S)*-equol-producing bacteria to produce DZN using DHD. Moreover, the final metabolite of DZN was DHD rather than *(S)*-equol for unknown reasons, and all of these results suggested that the metabolic pathway of *(S)*-equol production in Clostridium C1 can be regulated by other factors. To determine the new characteristics of *(S)*-equol production in Clostridium C1, whole genomes were sequenced, and *(S)*-equol-producing genes were identified by Blast comparison with reported functional genes. A gene named C1-07020 that showed 44.8% amino acid similarity with Lac 20–92 DZNR was obtained from bioinformatic analysis. Initially, the DZNR activity of C1-07020 was identified in whole-cell extracts using an *E. coli* expression system. In addition, another soy isoflavone, genistein, was also detected in this experiment, and this showed that genistein could be converted to dihydrogenistein (Fig. S7). Presumably, C1-07020 should be involved in the first step from DZN to *(S)*-equol or genistein to 5-hydroxyequol. Otherwise, there were no other genes with similarity to DDRC, DHDR and THDR that were found in Clostridium C1. It have been reported that all of the major known genes for *(S)*-equol production are located in a gene cluster with adjacent positions (23–27), and for our analyses of potential gene clusters in Clostridium C1, 30 genes located up and downstream of C1-07020 were investigated and some potential functional genes were also selected for enzyme activity detection *in vitro*, but no targets were found (Fig. S4). Therefore, Clostridium C1 may adopt a different mechanism to produce *(S)*-equol and other functional genes need to be further explored.

C1-07020 recombinant protein was purified and used for further functional assays *in vitro* in this work. A comparison of the DZN reductase activity between C1-07020 and DZNRs from Lac 20–92 and NATTS showed that all of these recombinant proteins possessed reductase activity for the conversion of DZN to DHD, and these enzyme reaction systems require NAD(P)H and strictly anaerobic conditions (39). Of which, NADH was the unique hydrogen donor for C1-07020. Remarkably, DZN could be converted to *(R)*-DHD using the known DZNRs, while only *(S)*-DHD can be metabolized to *(S)*-equol. DDRC was the key enzyme to affect the conversion of chiral molecules between *(R)*-DHD and *(S)*-DHD (23). However, about 70% of the product of *(S)*-DHD was detected in our C1-07020 enzyme reactions (Fig. 1A), meaning that C1-07020 has the activity of directly producing *(S)*-DHD without DDRC.

Interestingly, only C1-07020 was confirmed to have DHD oxidase activity that could reverse catalyze DHD to DZN by dehydrogenation, and was not affected by chiral molecules. This illustrated that C1-07020 was a novel bifunctional gene involved in producing *(S)*-equol. Moreover, the DHD reaction kinetic revealed that the enzyme reaction rate of C1-07020 was significantly lower than the rates of DZNRs, which could be relevant for its reversible enzyme activity. We also found that the optimal pH for DZN reductase activity (pH 7.5) and the optimal pH for DHD oxidase activity (pH 6.5) were different, and the direction of these reactions may be associated with the relative pH.

In our previous work, based on the four functional genes (*DZNR, DDRC, DHDR*, and *THDR*) from Lac 20–92, an engineering bacterium DDDT was successfully constructed and produced *(S)*-equol *in vitro*. To further evaluate the role of C1-07020, this gene was used to replace DZNR in the DDDT strain and the production of *(S)*-equol was compared by fermentation experiments. We found that C1-07020DDT exhibited higher *(S)*-equol production than DDDT (p < 0.05), although the fermenting time was longer, which corresponds to the enzyme kinetics of C1-07020 (Fig. 1B). Moreover, DZN was found in the fermentation of C1-07020DDT using *(S)*-equol as a substrate, and the solution pH exhibited an obvious impact on the formation between *(S)*-equol and DZN.

To be active as a DZN reductase, NAD(P)H and anaerobic conditions are essential. Indeed, the conversion ratio in *E. coli* was significantly higher than the specific activities of the purified enzymes for C1-07020 and DZNR (39). Conversion of purified enzymes was not improved even with NAD(P)H at high concentrations (Fig. S8). Specifically, C1-07020 still retained DHD oxidase activity without NADH or anaerobic conditions, although it could not transform DHD to DZN completely *in vitro* (Fig. 2). This indicated that some unknown cofactor may responsibility for the enzyme activity, and exhibited that the direction of the reaction was more favorable for DHD oxidases, which goes in the opposite direction of *(S)*-equol production in Clostridium C1. Accordingly, a possible reason for producing *(S)*-equol instability in this bacterium may be the direction of the enzyme reaction for C1-07020.

We also investigated the influence of *(S)*-equol and estradiol on the feedback enzyme activity of C1-07020, and our results indicated that *(S)*-equol could apparently promote DHD oxidase and inhibit DZN reductase activity at concentrations exceeding 0.64mM. Although *(S)*-equol is an estrogen-like chemical, estradiol did not exert similar effects in our hands (Fig. 4). These observations lead to the direct modulation of enzyme activity of C1-07020 by *(S)*-equol. Thus, a high concentration of *(S)*-equol may be a feedback negative regulator of this *(S)*-equol-producing pathway in Clostridium C1. In contrast, the DZN reductase activity of DZNRs was directly inhibited by *(S)*-equol (40). *(S)*-equol is known to be bacteriostatic to species such as *Clostridioides difficile*, Salmonella and some fungi (11, 12, 41), and the level of *(S)*-equol and relative bacteria can change the intestinal flora or representative bacterial species in human and animal guts, which is beneficial to the conditions for growth of these organisms (33, 42–44). However, high concentrations of *(S)*-equol can also suppress growth, and according to the reversibility between DZN and *(S)*-equol, it can be inferred that a feedback loop regulation may exist for the production of *(S)*-equol in Clostridium C1.

Bioinformatics analysis indicated that both C1-07020 and DZNRs have three conserved domains, an OYE-like flavin mononucleotide (FMN) binding domain, a 4Fe-4S cluster motif and a coenzyme binding motif (45). Hydride transfer from cofactors plays an important role in many redox reactions, and often involves cofactors such as flavin, deazaflavin and nicotinamide adenine dinucleotide (NAD) (45). Different reductases may have different coenzyme-binding sites involved in electron transfer, in which x amino acids from these three conserved domains may determine the selectivity of cofactors (Fig. S5). For the DZN reductase activity of C1-07020, NADH but not NADPH provided hydrogen ions in DZN reduction reactions.

To investigate the novel function of C1-07020, we first compared all DZNR genes with C1-07020. It could be seen that the most highly conserved domains were located in the N-terminus of amino acids 1–395, and the N-terminus had a higher sequence similarity than the C-terminus. The Swiss-model 3D structure of C1-07020 showed that the N-terminal and C-terminal remained relatively separated, which meant that the novel enzyme activity region may have been located in the C-terminus. Then, we utilized a truncation strategy with the C-terminal protein. A total of four truncated proteins, which were cut into regions from 371 to 643 aa, 531 to 643 aa, 584 to 643 aa, and 615 to 643 aa, were expressed in *E. coli* and identified *in vitro*, but a mere 28 amino acid truncation resulted in the loss of all enzyme activity, which meant that the C-terminus was important for the normal function of C1-07020 (Fig. S9).

Afterward, molecular docking and site-directed mutagenesis was performed and used for the evaluation of the binding sites in C1-07020 and DZN/DHD. We found that the residue Cys-72 was predicted to interact with the chromone scaffold of the DHD, but it had no effect on any enzyme activity. In contrast, neighboring residues, with Arg-75, were the key residues for the DHD oxidases activity of C1-07020 (Fig. 3). Moreover, we found that Lac 20–92 DZNR gained DHD oxidase activity when Ser-75 was mutated to Arg-75 (Fig. S6), which meant that Arg-75 may have been the key residue for DHD oxidase activity in DZNRs.

*(S)*-equol-producing bacteria are necessary for producing *(S)*-equol in human and animal intestines. However, many individuals who have these bacteria in their feces had undetectable levels of *(S)*-equol, even after supplementation with a soy diet (10, 31, 32, 34). Based on the *(S)*-equol-producing features and the discovery of a bifunctional gene in Clostridium C1, metagenomics analysis and PCR detection were used to evaluate the distribution of C1-07020-like genes in human and mouse intestines, respectively. We found more than 20 genes whose amino acid sequence similarity was higher than 80% to C1-07020 from human gut metagenomic datasets (Fig. 3). To our surprise, one gene and C1-07020 were completely identical, and another gene showing 80% similarity to C1-07020 was shown to have the same function as C1-07020 (Fig. 6A). Moreover, all genes in cluster 1 showed the same amino acid sequences in their functional sites as C1-07020, which indicated that these genes may have the same enzyme activity. Additionally, the reported DZNRs with lower sequence similarities but similar functions as C1-07020 belonged to cluster 2, but no DZNR was found when gene similarity with C1-07020 was lower than 40% (Fig. 6B).

It has been reported that all rodents, but not humans, have the ability of producing *(S)*-equol (10). In this study, only part of the genes in cluster 2 were detectable in mice. However, almost all of them were able to detect *(S)*-equol as well as the genes of cluster 1, but the detection rates showed no obvious correlation with *(S)*-equol yield (Fig. 6C). In humans, although the proportion of *(S)*-equol producers were very low (30%), the C1-07020-like genes were similar to mice (Fig. 6C), therefore, this novel *(S)*-equol-producing gene may be widespread across at least human, chick and mouse gut.

Overall, it could be deduced that Clostridium C1-like bacteria or genes with *(S)*-equol production roles may be commonly present in the intestine relative to known *(S)*-equol-producing bacteria. *(S)*-equol production capacity of an individual may not be dependent on the amount and species present in the gut, but rather tightly dependent on these colonies in a favorable intestinal environment. In these colonies, the bifunctional genes that are involved in *(S)*-equol production are important targets for the metabolic regulation of *(S)*-equol.

## ACKNOWLEDGMENTS

We thank LetPub (www.letpub.com) for its linguistic assistance during the preparation of this manuscript.

## REFERENCES

1. Ma L, Liu G, Ding M, Zong G, Hu FB, Willett WC, Rimm EB, Manson JE, Sun Q. 2020. Isoflavone intake and the risk of coronary heart disease in us men and women: results from 3 prospective cohort studies. Circulation 141:1–11.

2. Battaglia A, Devos G, Boeckx G, Goeman L, Tosco L, de Meerleer G, Joniau S. 2020. Prostate-specific antigen modulatory effect of a fermented soy supplement for patients with an elevated risk of prostate cancer: a non-randomized, retrospective observational registration. Curr Urol 14:142–149.

3. Katagiri R, Sawada N, Goto A, Yamaji T, Iwasaki M, Noda M, Iso H, Tsugane S. 2020. Association of soy and fermented soy product intake with total and cause specific mortality: prospective cohort study. The BMJ 368:m34.

4. Matsumoto T, Kojima M, Takayanagi K, Taguchi K, Kobayashi T. 2020. Role of S-equol, indoxyl sulfate, and trimethylamine n-oxide on vascular function. Am J Hypertens 33:793–803.

5. Zheng W, Ma Y, Zhao A, He TH, Lyu N, Pan ZQ, Mao GQ, Liu Y, Li J, Wang PY, Wang J, Zhu BL, Zhang YM. 2019. Compositional and functional differences in human gut microbiome with respect to equol production and its association with blood lipid level: a cross-sectional study. Gut Pathog 11:20.

6. Setchell KD, Clerici C. 2010. Equol: pharmacokinetics and biological actions. J Nutr 140:1363S–8S.

7. Ahuja V, Miura K, Vishnu A, Fujiyoshi A, Evans R, Zaid M, Miyagawa N, Hisamatsu T, Kadota A, Okamura T, Ueshima H, Sekikawa A. 2017. Significant inverse association of equol-producer status with coronary artery calcification but not dietary isoflavones in healthy Japanese men. Br J Nutr 117:260–266.

8. Yoshikata R, Myint KZ, Ohta H. 2017. Relationship between equol producer status and metabolic parameters in 743 Japanese women: equol producer status is associated with antiatherosclerotic conditions in women around menopause and early postmenopause. Menopause 24:216–224.

9. Lambert M, Thybo CB, Lykkeboe S, Rasmussen LM, Frette X, Christensen LP, Jeppesen PB. 2017. Combined bioavailable isoflavones and probiotics improve bone status and estrogen metabolism in postmenopausal osteopenic women: a randomized controlled trial. Am J Clin Nutr 106:909–920.

10. Mayo B, Vazquez L, Florez AB. 2019. Equol: a bacterial metabolite from the daidzein isoflavone and its presumed beneficial health effects. Nutrients 11:1–20.

11. Tanaka Y, Kimura S, Ishii Y, Tateda K. 2019. Equol inhibits growth and spore formation of *Clostridioides difficile*. J Appl Microbiol 127:932–940.

12. Li BU. 2019. Advances in exploring equol production and application. J Food Process Pres 43:e14205.

13. Chen LR, Ko NY, Chen KH. 2019. Isoflavone supplements for menopausal women: a systematic review. Nutrients 11:2649.

14. Yoshikata R, Myint K, Ohta H, Ishigaki Y. 2021. Effects of an equol-containing supplement on advanced glycation end products, visceral fat and climacteric symptoms in postmenopausal women: A randomized controlled trial. PLoS One 16:e0257332.

15. Kurylowicz A. 2020. The role of isoflavones in type 2 diabetes prevention and treatment-a narrative review. Int J Mol Sci 22.

16. Daily JW, Ko BS, Ryuk J, Liu M, Zhang W, Park S. 2019. Equol decreases hot flashes in postmenopausal women: a systematic review and Meta-analysis of randomized clinical trials. J Med Food 22:127–139.

17. Kwon JE, Lim J, Bang I, Kim I, Kim D, Kang SC. 2019. Fermentation product with new equol-producing *Lactobacillus paracasei* as a probiotic like product candidate for prevention of skin and intestinal disorder. J Sci Food Agr 99:4200–4210.

18. Dai S, Pan M, El-Nezami HS, Wan J, Wang MF, Habimana O, Lee J, Louie J, Shah NP. 2019. Effects of lactic acid bacteria-fermented soymilk on isoflavone metabolites and short-chain fatty acids excretion and their modulating effects on gut microbiota. J Food Sci 84:1854–1863.

19. Feng XL, Ho SC, Zhan XX, Zuo LS, Mo XF, Zhang X, Abulimiti A, Huang CY, Zhang CX. 2021. Serum isoflavones and lignans and odds of breast cancer in pre-and postmenopausal Chinese women. Menopause 28:413–422.

20. Lu C, Lv JW, Jiang N, Wang HX, Huang H, Zhang LJ, Li SY, Zhang NN, Fan B, Liu XM, Wang FZ. 2020. Protective effects of genistein on the cognitive deficits induced by chronic sleep deprivation. Phytother Res 34:1–13.

21. Cho HW, Gim HJ, Li H, Subedi L, Kim SY, Ryu JH, Jeon R. 2021. Structure-activity relationship of phytoestrogen analogs as eralpha/beta agonists with neuroprotective activities. Chem Pharm Bull (Tokyo) 69:99–105.

22. Setchell KD, Cole SJ. 2006. Method of defining equol-producer status and its frequency among vegetarians. J Nutr 136:2188–93.

23. Shimada Y, Takahashi M, Miyazawa N, Abiru Y, Uchiyama S, Hishigaki H. 2012. Identification of a novel dihydrodaidzein racemase essential for biosynthesis of equol from daidzein in *Lactococcus sp.* strain 20-92. Appl Environ Microbiol 78:4902–7.

24. Schroder C, Matthies A, Engst W, Blaut M, Braune A. 2013. Identification and expression of genes involved in the conversion of daidzein and genistein by the equol-forming bacterium *Slackia isoflavoniconvertens*. Applied and environmental microbiology 79:3494–502.

25. Tsuji H, Moriyama K, Nomoto K, Akaza H. 2012. Identification of an enzyme system for daidzein-to-equol conversion in *Slackia* sp. strain NATTS. Appl Environ Microbiol 78:1228–36.

26. Kawada Y, Yokoyama S, Yanase E, Niwa T, Suzuki T. 2016. The production of S-equol from daidzein is associated with a cluster of three genes in *Eggerthella* sp. YY7918. Biosci Microbiota Food Health 35:113–21.

27. Florez AB, Vazquez L, Rodriguez J, Redruello B, Mayo B. 2019. Transcriptional regulation of the equol biosynthesis gene cluster in *Adlercreutzia equolifaciens* DSM19450(T). Nutrients 11:993.

28. Li HL, Mao SM, Chen HH, Zhu LY, Liu W, Wang X, Yin YS. 2018. To construct an engineered (S)-equol resistant E. coli for in vitro (S)-equol production. Front Microbiol 9:1182.

29. Lee PG, Lee SH, Kim J, Kim EJ, Choi KY, Kim BG. 2018. Polymeric solvent engineering for gram/liter scale production of a water-insoluble isoflavone derivative, (S)-equol. Appl Microbiol Biotechnol 102:6915–6921.

30. Sajid M, Stone SR, Kaur P. 2021. Recent advances in heterologous synthesis paving way for future green-modular bioindustries: a review with special reference to isoflavonoids. Front Bioeng Biotechnol 9:673270.

31. Iino C, Shimoyama T, Iino K, Yokoyama Y, Chinda D, Sakuraba H, Fukuda S, Nakaji S. 2019. Daidzein intake is associated with equol producing status through an increase in the intestinal bacteria responsible for equol production. Nutrients 11:433.

32. Yoshikata R, Myint KZ, Ohta H, Ishigaki Y. 2019. Inter-relationship between diet, lifestyle habits, gut microflora, and the equol-producer phenotype: baseline findings from a placebo-controlled intervention trial. Menopause 26:273–285.

33. Vazquez L, Florez AB, Guadamuro L, Mayo B. 2017. Effect of soy isoflavones on growth of representative bacterial species from the human gut. Nutrients 9:727.

34. Vazquez L, Guadamuro L, Giganto F, Mayo B, Florez AB. 2017. Development and use of a real-time quantitative PCR method for detecting and quantifying equol-producing bacteria in human faecal samples and slurry cultures. Front Microbiol 8:1155.

35. Trott O, Olson AJ. 2010. AutoDock Vina: improving the speed and accuracy of docking with a new scoring function, efficient optimization, and multithreading.J Comput Chem 31:455–61.

36. Goodsell DS, Sanner MF, Olson AJ, Forli S. 2021. The AutoDock suite at 30. Protein Sci 30:31–43.

37. Almeida A, Nayfach S, Boland M, Strozzi F, Beracochea M, Shi ZJ, Pollard KS, Sakharova E, Parks DH, Hugenholtz P, Segata N, Kyrpides NC, Finn RD. 2021. A unified catalog of 204,938 reference genomes from the human gut microbiome. Nat Biotechnol 39:105–114.

38. Tamura K, Stecher G, Kumar S. 2021. MEGA11: molecular evolutionary genetics analysis version 11. Mol Biol Evol 38:3022–3027.

39. Shimada Y, Yasuda S, Takahashi M, Hayashi T, Miyazawa N, Sato I, Abiru Y, Uchiyama S, Hishigaki H. 2010. Cloning and expression of a novel NADP(H)-dependent daidzein reductase, an enzyme involved in the metabolism of daidzein, from equol-producing *Lactococcus* strain 20-92. Appl Environ Microbiol 76:5892–5901.

40. Lee PG, Kim J, Kim EJ, Jung E, Pandey BP, Kim BG. 2016. P212A mutant of dihydrodaidzein reductase enhances (s)-equol production and enantioselectivity in a recombinant *Escherichia coli* whole-cell reaction system. Appl Environ Microbiol 82:1992–2002.

41. Wang JY, Li L, Yin YS, Gu ZK, Chai RY, Wang YL, Sun GC. 2017. Equol, a clinically important metabolite, inhibits the development and pathogenicity of Magnaporthe oryzae, the causal agent of rice blast disease. Molecules 22:1799.

42. Bolca S, Verstraete W. 2010. Microbial equol production attenuates colonic methanogenesis and sulphidogenesis in vitro. Anaerobe 16:247–52.

43. Zheng WJ, Hou YJ, Su Y, Yao W. 2014. Lactulose promotes equol production and changes the microbial community during in vitro fermentation of daidzein by fecal inocula of sows. Anaerobe 25:47–52.

44. Zheng WJ, Zhang X, Yao W. 2016. Individual difference in faecal and urine equol excretion and their correlation with intestinal microbiota in large white sows. Anim Prod Sci 57:262–270.

45. Kawada Y, Goshima T, Sawamura R, Yokoyama SI, Yanase E, Niwa T, Ebihara A, Inagaki M, Yamaguchi K, Kuwata K, Kato Y, Sakurada O, Suzuki T. 2018. Daidzein reductase of *Eggerthella* sp. YY7918, its octameric subunit structure containing FMN/FAD/4Fe-4S, and its enantioselective production of R-dihydroisoflavones. J Biosci Bioeng 126:301–309.

